# Turgor pressure affects transverse stiffness and resonant frequencies of buzz-pollinated poricidal anthers

**DOI:** 10.1101/2024.05.21.595227

**Authors:** Mitchell Alvord, Jenna McNally, Cailin Casey, Mark Jankauski

## Abstract

Several agriculturally valuable plants store their pollen in tube-like poricidal anthers, which release pollen through buzz pollination. In this process, bees rapidly vibrate the anther using their indirect flight muscles. The stiffness and resonant frequency of the anther are crucial for effective pollen release, yet the impact of turgor pressure on these properties is not well understood. Here, we perform three-point flexure tests and experimental modal analysis to determine anther transverse stiffness and resonant frequency, respectively. Dynamic nanoindentation is used to identify the anther’s storage modulus as a function of excitation frequency. We subsequently develop mathematical models to estimate how turgor pressure changes after the anther is removed from a flower, thereby emulating zero water availability. We find that 30 minutes post-ablation, anther stiffness decreases 60%. Anther resonant frequency decreases 20% 60 minutes post-ablation. Models indicate turgor pressure in the fresh anther is about 0.2 – 0.3 MPa. Our findings suggest that natural fluctuations in turgor pressure due to environmental factors like temperature and light intensity may require bees to adjust their foraging behaviors. Interestingly, anther storage modulus increased with excitation frequency, underscoring the need for more sophisticated mechanical models that consider viscous fluid transport through plant tissue moving forward.

## Introduction

Approximately 10% of flowering plants hold pollen in tube-like structures called poricidal anthers [1, 2, 3]. Pollen is extracted from poricidal anthers via ‘buzz pollination’, a behavior performed by bumblebees, carpenter bees, mason bees and other bee species [4, 5, 6]. During buzz pollination, a bee will bite an anther with its mandibles and rapidly vibrate the anther by contracting its indirect flight muscles [7, 8]. Vibration causes the pollen to exit the anther’s apical pores where it is collected on the bee’s abdomen. The pollen is then transferred to other flowers as the bee continues to forage. This process is essential for the reproduction of several agriculturally valuable crops, including tomatoes, eggplant, blueberries, and so on [9]. Although the evolution and ecology of buzz pollination have been studied extensively [3, 10, 11, 12, 13, 14, 15, 16, 17, 18, 19], the mechanics of buzz-pollinated poricidal anthers have received comparatively less attention (but see [20, 21, 22, 23, 24, 25]).

The success of pollen extraction during buzz pollination depends on the mechanics of the flexible anther. The compliant anther bends under the forces applied by the bee, thereby causing the anther’s interior locule walls to vibrate and dislodge pollen grains [26]. The amount of pollen released by the anther is correlated with the anther’s vibration amplitude [27]. Anther vibration amplitude is influenced by several factors, including the magnitude and frequency of the forces applied by the bee and the location where the bee bites the anther [28]. Bees appear to take advantage of vibration amplification to increase the deformation of the anther [29]. Vibration amplification usually involves exciting a structure harmonically near its resonant frequency, where the resonant frequency refers to the specific frequency at which an object naturally vibrates or oscillates most prominently when energy is applied [30]. When exciting a structure near resonance, small periodic forces elicit large structural responses. Anther resonant frequencies are governed by their mass distribution and transverse stiffness, where the transverse stiffness is the resistance to deflection or bending in a direction perpendicular to its longitudinal axis. Practically, anther resonant frequencies are a function of morphological and material properties. However, transverse stiffness and resonant frequencies may also be influenced by turgor pressure, yet the role that turgor pressure plays in the mechanics of poricidal anthers has not been adequately addressed.

Turgor pressure is a hydrostatic pressure that occurs in plants due to the osmotic flow of water that pushes the cell membrane against the cell wall [31]. Methods have been developed to measure the turgor pressure on the organ scale, such as the pressure bomb [32], as well as on the tissue or cellular scale, such as pressure probe or indentation techniques [33]. It is well established that turgor pressure affects the rigidity of plant structures; reductions in turgor pressure are associated with wilting [34]. Several factors influence the turgor pressure within plant tissues, including water availability, temperature, and soil conditions [35, 36, 37]. Therefore, the turgor pressure is a dynamic property that may change in response to environmental conditions. This implies that anther ridigity and resonant frequencies may also vary with changes in the environment. Consequently, if bees do leverage resonance or vibration amplification as previously hypothesized, they must adapt to the shifts in anther resonant frequencies caused by changes in turgor pressure. This may drive changes in behavior, such as modulating their buzzing frequencies to coincide with the anther structural resonance.

Here, we investigate the influence of turgor pressure on the transverse stiffness and first resonant frequency (simply resonant frequency hereafter) of poricidal anthers of *Solanum sisymbriifolium* flowers. Rapid loss of turgor pressure was induced by removing the anther from the flower and depriving the filament of water. Although organ-level methods have been developed to measure turgor pressure directly [33], these methods are often invasive and are not conducive to the time-resolved measurements required in the present study. We therefore developed a novel approach to estimate the turgor pressure based on mechanical modeling and structure-level experiments. We experimentally measured anther transverse stiffness using a three-point flexure test and the resonant frequency through experimental modal analysis. Additionally, we performed dynamic indentation to characterize anther tissue storage modulus as a function of frequency, and conducted a simple morphological analysis to measure anther length and effective radius. We then compared the measured anther transverse stiffness and resonant frequency to those predicted by mathematical models. We hypothesized that reductions in turgor pressure would lower the anther’s transverse stiffness and resonant frequencies.

## Materials and Methods

### Experimental Procedures

#### Specimen Collection and Morphometrics

We ordered *S. sisymbriifolium* plants in the vegetative state to Bozeman, Montana from the Sow Exotic Nursery (Winter Haven, FL, USA). Plants were grown at the Montana State University Plant Growth Center until they began to flower. We cut fresh flowers at the stem and placed them in a centrifugal tube with water to prevent withering. The flowers were cut and tested within the same day.

We quantified the length and diameters of *S. sysimbriifolium* anthers (*n* = 14) by conducting a morphometric study. We removed the anthers from the flower at the filament, weighed them, and imaged them on grid paper. We used ImageJ to measure the effective diameter and length of the anther. Due to the anther’s variable cross-section, we measured the diameter at five different points equally spaced along the anther length and averaged them to obtain a representative diameter.

#### Three Point Flexure Test

We performed a quasi-static three-point flexure test on anthers (n = 5) to measure their transverse stiffness over time after the anther was ablated from the flower (Fig. 1). A quasi-static test was required because the anthers exhibited relaxation behavior when deformed with a purely static load.

**Fig. 1.**
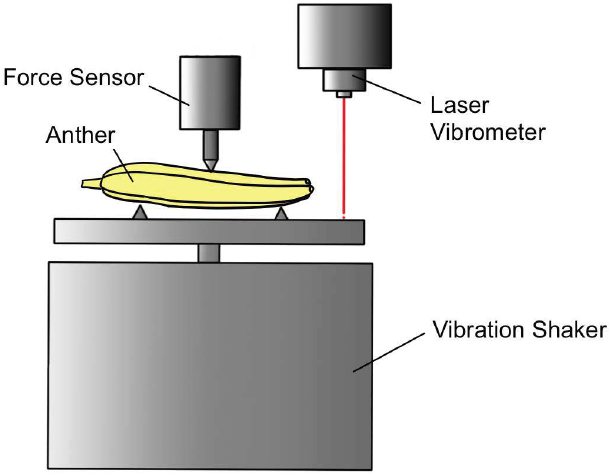
Experimental setup used to perform three-point flexure test.

**Fig. 2.**
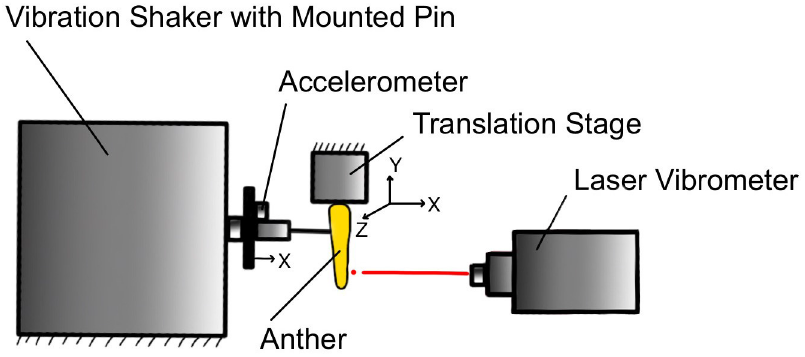
Experimental setup used to perform experimental modal analysis.

To perform the three-point flexure test, we placed the anther on a 3D printed fixture with two triangular prisms with a pin-to-pin length of 5 mm. The fixture was connected to an electrodynamic shaker (2007E, The Modal Shop, Cincinnati, OH, USA) with an external amplifier (2100E21-100, The Modal Shop, Cincinnati, OH, USA). Shaker displacement was measured via a laser vibrometer (Polytec VGO - 200, Polytec, Baden-Württemberg, Germany) with a sensitivity of 250 *μ*m V^−1^. We used a load cell (GS0-10, Transducer Techniques, Temecula, CA, USA) to measure the corresponding force due to the deflection of the anther. The load cell was positioned to the midpoint of the anther between the pins. The shaker motion followed a 10 Hz rectified sine wave and an amplitude of approximately 100 *μ*m. The rectified signal was chosen such that during testing the load cell remained in contact with the anther. Data were recorded using a data acquisition system (cDAQ 9178, National Instruments, Austin, TX, USA) with a sampling frequency of 1000 Hz.

We performed the initial test on the anther approximately three minutes after removal from the flower. Subsequent measurements occurred starting at five minutes after removal and then in five-minute increments up to thirty minutes. We determined the transverse stiffness by plotting the force vs. deflection curve and taking a linear regression. The slope of the regression line defined the transverse stiffness of the anther.

#### Experimental Modal Analysis

We performed experimental modal analysis to determine how the anther’s resonant frequency was affected by pressure loss (Fig. **??**). Tests were performed on *n* = 5 anthers. The anther base was secured to a 3D printed cube with a centrally located hole via hot glue. We then placed the anther in an inverted position into a clamp connected to a translation stage. We applied a force directly to the anther using an electrodynamic shaker (K2007E01, The Modal Shop, Cincinnati, OH, USA), where the anther and shaker were connected via a narrow gauge insect pin. The insect pin was located approximately 30% of the anther length from the base, near where a bee bites the anther during buzz pollination. The pin-anther interface was secured via a small bead of glue. Measurements began 10 minutes after the anther was ablated from the flower and continued in 5-minute increments up to 60 minutes after detachment.

The shaker was prescribed a periodic chirp signal over a frequency range of 0.2 - 5 kHz lasting 640 milliseconds. A total of 15 complex averages were used to compute the fast Fourier transform of the time series acceleration and velocity data. The duration of each test was 10 to 15 seconds. The input acceleration at the pin location was measured using an accelerometer (352A21, PCB Piezotronics, Depew, NY, USA) with a sensitivity of 10.2 mV g^−1^. The anther response velocity was measured via a laser vibrometer (PSV-400, Polytec, Hudson, MA, USA) with close-up attachment (PSV-A-410, Polytec, Hudson, MA, USA) at a sensitivity of 50 mm s^−1^ V^−1^. Input shaker accelerations ranged from 3 to 10 m s^−2^ and anther velocities ranged from 0.5 to 40 mm s^−1^ over the duration of one chirp. The deformation velocity of the anther and reference acceleration of the shaker were then reconstructed into frequency response functions using Polytec Presentation Viewer. The resonant frequency was extracted from the frequency response using a local curve fitter within the Polytec software.

#### Dynamic Nanoindentation

We performed dynamic nanoindentation to identify the storage modulus of the poricidal anther as a function of excitation frequency. Poroelastic tissues, such as plant tissue, are imbued with water. At high excitation frequencies, the water may not have sufficient time to flow between adjoining pores, thereby increasing the effective storage modulus. Therefore, it is appropriate to use different moduli values for the mathematical models describing the quasi-static flexure test, which occurs at low frequency, and the experimental modal analysis dynamic test, which spans a higher frequency range.

First, the anther was cut in the longitudinal direction. The cut anther was then embedded in slow cure epoxy resin (Pelco, Ted Pella, Redding, CA, USA) for at least a week to allow the epoxy to reach full strength. Slow-curing epoxy resin does not require heat and has a high level of penetration to properly embed the intricate poroelastic tissue. Embedded samples were cut in the transverse plane to expose a cross-section of the anther using a slow speed saw with a diamond blade. The sample surface was polished first using 600 grit and 1000 grit sand paper and then in a series of liquid diamond polishing solution from 9μm to 0.05 μm (Ted Pella, Redding, CA, USA). The sample was submerged in water to maintain hydration during testing. Four indents were placed on the anther with a spacing of 100 μm, and maximum indentation depth was about 1200 nm. Dynamic nanoindentation was performed using the iMicro nanoindenter (KLA Tencor, Milpitas, California, USA) using a flat punch tip with a radius of 54.5 μm to measure storage modulus. Ten test frequencies were linearly spaced from 1 to 200 Hz.

### Mathematical Modeling

The anther turgor pressure was not measured directly within our experimental efforts. Conventional methods for measuring turgor pressure, such as the pressure bomb, are not conducive to monitoring how the turgor pressure changes over time. We instead rely on mathematical modeling coupled with experimental measurements to estimate how the turgor pressure varies after the anther is ablated from the flower. We first derived a static model that predicts the turgor pressure as a function of transverse stiffness. We then derive a dynamic model that predicts the turgor pressure as a function of the first resonant frequency. Both models require anther morphology and material properties. Here, we discuss the derivation and assumptions employed by each model.

#### Cross Section Geometry and Porosity

The cross section of the anther is complex and heterogeneous (Fig. 3). To simplify the modeling, we identified a single representative cross section that accounts for the four pollen cavities and plant tissue porosity.

**Fig. 3.**
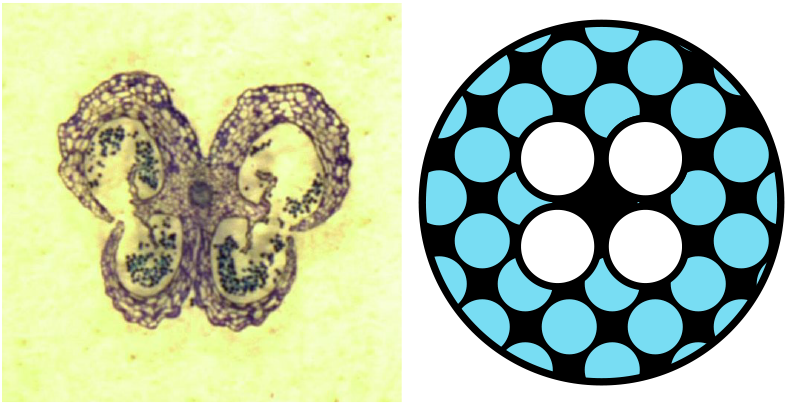
(Left) Histology slide of *S. sisymbriifolium* anther. (Right) Simplified cross section of poricidal anther. Four white circles represent the pollen cavities with radius *r*_*in*_. Blue areas represent the porous structure within the cross section. Black areas represent solid plant material.

The anther cross section was assumed circular with radius *r*_*o*_, where the outer radius was determined from morphological measurements. Based on histology slides, we assumed that the inner pollen cavities accounted for approximately 50% of the total anther cross sectional area. We idealized the four pollen cavities as circular and arranged them around the center of the anther’s longitudinal axis. Then, the radius of a single pollen cavity *r*_*in*_ is

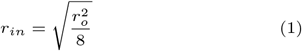

The cross sectional area of the anther can then be determined as the outer circle less the four pollen cavities. However, plant tissue is highly porous; we must therefore adjust the effective cross sectional area to account for this porosity, as the porous regions of the cross section will not affect the solid mechanical phase. We introduce a parameter *β* to denote porosity, where *β* is defined as the ratio of pore area to the overall cross sectional area. Thus, *β* = 0 denotes a solid cross section, while *β* = 1 denotes a hollow cross section. Taking into account porosity, the effective solid cross-sectional area *A*_*eff*_ and the effective cross sectional area of the pores *A*_*pore*_ are

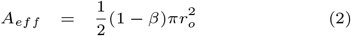

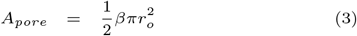

Lastly, using the same geometric and porosity assumptions, we approximate the second area of moment *I*_*eff*_ as

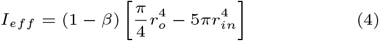

#### Static Modeling

We developed a static model of the poricidal anther to estimate the pressure from experimental measurements of transverse stiffness (Fig. 4). The anther was treated as a simply supported beam subject to an axial tensile load, where the tensile load represents the influence of turgor pressure on anther mechanics. The beam was broken into two segments of equal length *L*. The positions along the left and right segments are *x*_1_ and *x*_2_, respectively, and the deflection of the left and right segments are *W*_1_ and *W*_2_. The tensile load *T* imparted by the turgor pressure *p*_*t*_ is *T* = *p*_*t*_*A*_*pore*_. Assuming identical material and geometric properties between the two segments, the differential equation governing the deflection of the *i*^*th*^ beam segment is

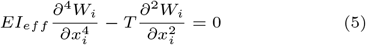

where *E* is the Young’s modulus and *I*_*eff*_ is the effective second area moment. The boundary conditions are identical at *x*_1_ = *x*_2_ = 0 and reflect no transverse displacement and no bending moment. The boundary conditions at *x*_1_ = *x*_2_ = *L* enforce kinematic compatability of displacement and rotation, and the equilibrium of bending moments. The total shear force in the beam at the geometric center is equivalent to the applied static force *F*. Under these assumptions, the anther’s transverse stiffness *k* and midpoint displacement *W*_*max*_ satisfies

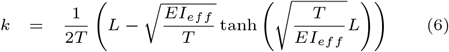

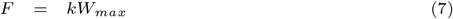

**Fig. 4.**
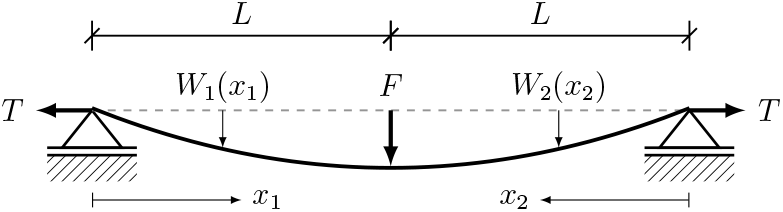
Model of an anther as a simply supported beam with axial load. The model is used to estimate turgor pressure based on experimental data.

#### Dynamic Modeling

Next, we develop a dynamic model to predict the natural frequencies of the poricidal anther with given geometry, material properties and turgor pressure (Fig. 5). Eq. 5 was modified to include the inertial force associated with anther transverse acceleration as

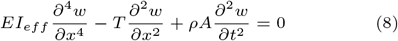

where *ρ* is the average material density of the anther’s solid and fluid phases and *A* is the total cross-sectional area. It can be shown that the vibration modes of the tensioned beam, which depend only on space, are described by [38]:

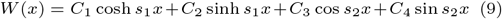

where *C*_1_−*C*_4_ are constants determined by boundary conditions and *s*_1_ and *s*_2_ are the roots of the auxiliary equation given by

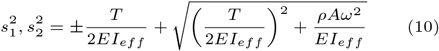

**Fig. 5.**
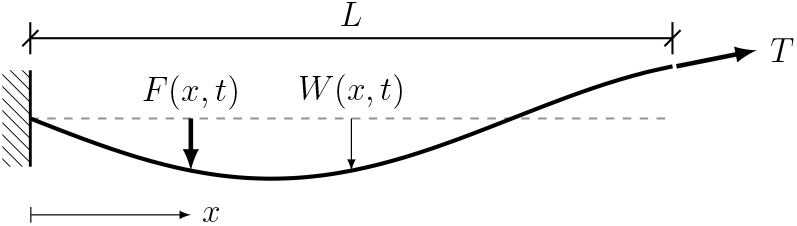
Mathematical model of an anther as a cantilever beam. The model is used to estimate the anther’s resonant frequency.

Here, *ω* is the oscillation frequency. To solve for the natural frequencies, we assume the beam is clamped at *x* = 0 such that it experiences no transverse deflection or rotation. The beam on the right edge where *x* = *L* experiences no bending moment, and the shear force is equal to the projection of *T* in the vertical direction.

Applying the boundary conditions to Eq. 9, we obtain a coefficient matrix that is dependent on *s*_1_ and *s*_2_. For *ω* to correspond to a resonant frequency *ω*_*n*_, the determinant of the coefficient matrix must be zero. Because no closed-form analytic solution for *ω*_*n*_ is possible, we numerically sweep through a range of frequencies and identify at which frequencies the determinant of the coefficient matrix becomes zero.

## Results

### Experimental Results

The results of the three-point flexure test are shown in Fig. 6. The anthers exhibited maximal transverse stiffness of about 385 N*·*m^−1^ on average immediately upon removal from the flower (*t* = 3 minutes). Transverse stiffness subsequently declined, falling by approximately 60% within 30 minutes after removal from the flower. At 30 minutes, the anthers exhibited small localized deformations in addition to large scale bending. Localized deformations occur at the pin locations referenced in Fig. 1. Due to these local deformations, it is likely that the transverse stiffness is underestimated at greater post-ablation times, where the turgor pressure is presumed to be lower.

**Fig. 6.**
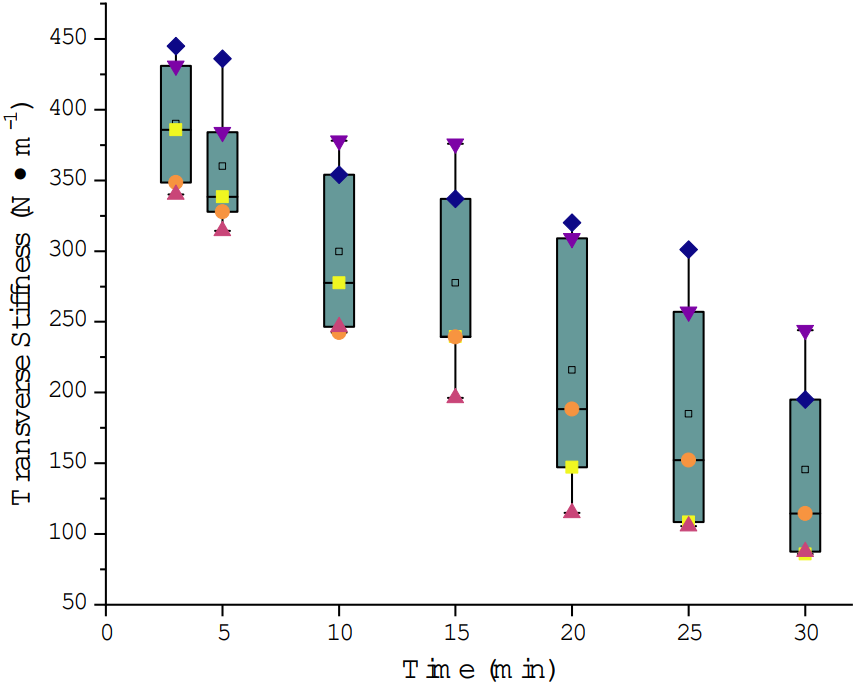
Anther transverse stiffness versus time for *n* = 5 anthers. Anther transverse stiffness was calculated from a 3-point flexure test.

The results of the experimental modal analysis test are shown in Fig. 7. The resonant frequency at *t* = 10 minutes occurs at about 625 Hz on average and declines to about 490 Hz at *t* = 60 minutes. The rate at which the resonant frequency changed was highest immediately after the anther was removed from the flower and tended to stabilize around 50 minutes post-ablation. The reduction in resonant frequency was more gradual compared to the reduction in transverse stiffness measured in the static flexure test.

**Fig. 7.**
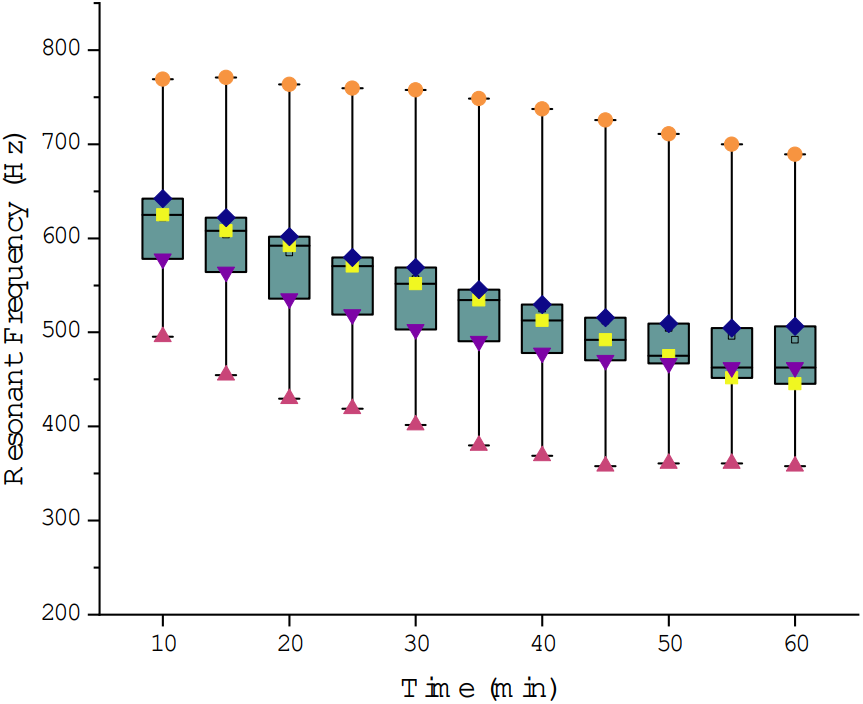
Anther resonant frequency versus time for *n* = 5 anthers. Anther resonant frequency was calculated from an experimental modal analysis test.

The results of the dynamic nanoindentation are shown in Fig. 8. The storage modulus was highly frequency-dependent; the average storage modulus considering static indentation was 18.4 MPa, compared to approximately 35.4 MPa with a dynamic indentation frequency of 200 Hz. The storage modulus increases rapidly at low frequencies and reaches a near steady value at about 100 Hz.

**Fig. 8.**
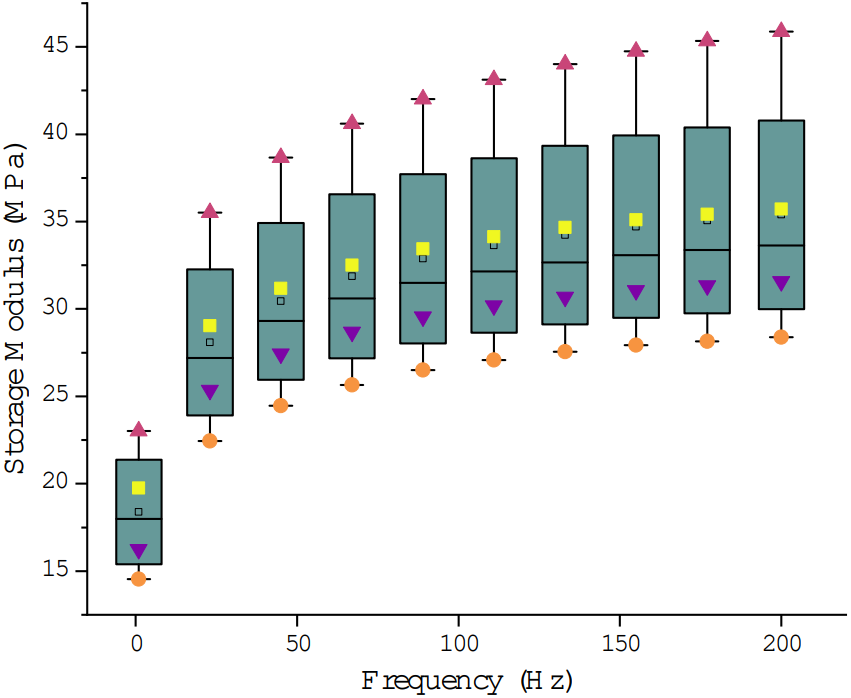
Anther storage modulus versus frequency for *n* = 1 anthers. Storage modulus was calculated from a dynamic nanoindentation test performed at four locations on an anther cross section.

### Model Results

The parameters used to populate the models are shown in Tab. 1. For *n* = 14 samples, the mean (± standard deviation) for outer anther radius was 0.8150 ± 0.068 mm, anther length was 8.6 ± 0.33 mm, and anther mass was 9.8 ± 1.5 mg, which yields an approximate density of 1000 kg *·* m^−3^. We used different modulus values for the static and dynamic models due to the frequency dependence of the storage modulus observed during dynamic nanoindentation (Fig. 8). We vary porosity *β* from 0.5 - 1.0 in 0.1 intervals, which is the approximate range for *β* where model predictions coincide with experimental measurements. We use static and dynamic models to solve for the transverse stiffness and first natural frequency, respectively, over a range of turgor pressures spanning 0 to 0.5 MPa in 100 evenly spaced increments.

**Table 1.** Parameters for static and dynamic anther models.

| Variable | Description | Value | Unit |
| --- | --- | --- | --- |
| $r_o$ | Effective Anther Radius | 0.8150 | mm |
| $\rho$ | Mass Density | 1000 | $\text{kg} \cdot \text{m}^{-3}$ |
| $E_s$ | Young's Modulus, Static | 18.4 | MPa |
| $E_d$ | Young's Modulus, Dynamic | 35.4 | MPa |
| $L_s$ | Static Test Length | 5 | mm |
| $L_d$ | Dynamic Test Length | 8.6 | mm |
| $\beta$ | Porosity | 0.5 - 1.0 | - |

The results of the static model are shown in Fig. 9. In all cases, the transverse stiffness decreases monotonically with turgor pressure. The rate at which the transverse stiffness decreases with pressure is the highest in the most porous case (*β* = 1), where the transverse stiffness is derived exclusively from the tensile force acting on the surface. In this case, the anther mathematically behaves as a tensioned string. By contrast, the rate at which transverse stiffness decreases with pressure is considerably less when *β* = 0.5, where a large percentage of transverse stiffness stems from the flexural rigidity of the anther’s solid phase. The dynamic model shows that turgor pressure has a similar influence on the anther’s resonant frequency as it does transverse stiffness (Fig. 10). The resonant frequency decreases monotonically with turgor pressure as well, and lower porosity values correspond to higher resonant frequencies throughout the range of turgor pressures considered. At high turgor pressures, the anther behaves as a tensioned string, while at low turgor pressures, it behaves more as a beam.

**Fig. 9.**
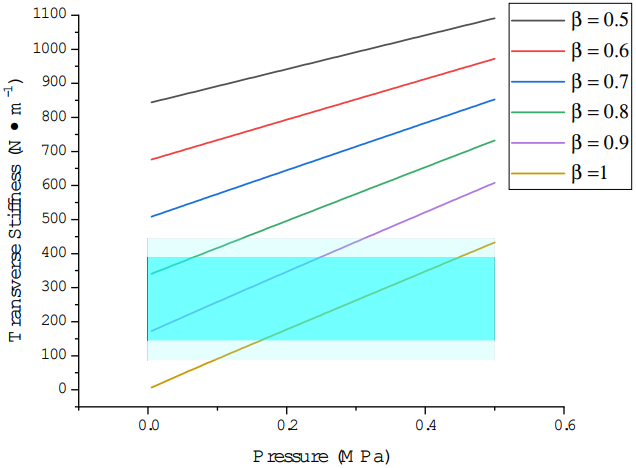
Turgor pressure versus transverse stiffness for different porosities *β* as predicted by static modeling. The light cyan box indicates the entire experimental range of stiffness values measured over time and the dark cyan box indicates the mean range of stiffness values measured over time.

**Fig. 10.**
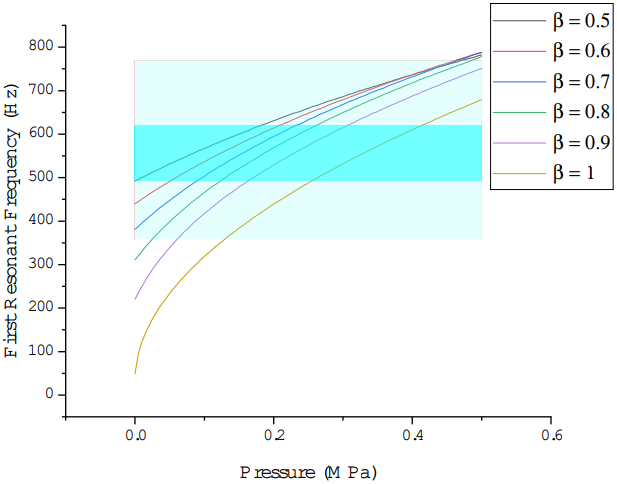
Turgor pressure versus resonant frequencies for different porosities *β* as predicted by dynamic modeling. The light cyan box indicates the entire experimental range of stiffness values measured over time and the dark cyan box indicates the mean range of stiffness values measured over time.

Both static and dynamic models can be used to approximate turgor pressure of the anthers considered in the 3-point flexure and experimental modal tests. In Figs. 9-10, the light cyan region denotes the total range of experimentally measured stiffness or resonant frequencies across all anthers and times, and the dark cyan box indicates the range of mean values at each time. For the static model, the anther must have relatively high porosity values of about *β* = 0.8 - 1.0 for the model to coincide with the experimental range. For porosities lower than *β* = 0.8, the stiffness at zero turgor pressure exceeds that of the mean experimental range at *t* = 30 minutes. For a porosity of *β* = 1.0, the static model predicts the turgor pressure in a freshly ablated anther with a transverse stiffness 375 N m^−1^ is about 0.45 MPa; for *β* = 0.9, the turgor pressure is about 0.25 MPa.

In contrast to the static model, the dynamic model predicts resonant frequencies in the mean experimental range over a wider span of porosities and pressures. For *β* = 1.0, the dynamic model predicts turgor pressures of about 0.45 MPa and 0.25 MPa at *t* = 10 minutes (resonant frequency of 625 Hz) and *t* = 60 minutes resonant frequency of 500 Hz) post-ablation, respectively. If *β* = 0.5, these values fall to 0.2 MPa and 0 MPa at *t* = 10 minutes and *t* = 60 minutes post-ablation, respectively.

To our knowledge, the porosity of any poricidal anther, including *S. sisymbriifolium* anthers, has not been measured. However, porosity has been measured at approximately 0.1 - 0.6 in other plant tissues [39, 40]. Based on our models, we hypothesize that the porosity of the *S. sisymbriifolium* anthers is the range of 0.5 - 0.9, and is more likely on the lower end. This corresponds to turgor pressures in freshly ablated anthers of 0.2 - 0.3 MPa, which is similar in magnitude to the turgor pressure measured in other plant tissues and structures [33]. The range of porosities predicted by the dynamic model is likely more accurate than the limited range of porosities predicted by the static model. This stems from the underestimation of the transverse stiffness due to local deformations at the pin locations in the three-point flexure test. However, results from both models are sensitive to changes in effective anther radius *r*_*o*_, since the second area moment is proportional to the fourth power of *r*_*o*_. Therefore, small changes in *r*_*o*_ will considerably affect transverse stiffness and resonant frequencies. Further, other assumptions such as uniform geometric and material properties may limit the absolute accuracy of modeling efforts. Still, both models agree with experimental trends and are useful tools in estimating how turgor pressure affects anther mechanics.

## Discussion

The mechanics of buzz-pollinated poricidal anthers are influenced by anther morphology, material properties, and turgor pressure. However, the role that turgor pressure plays in anther mechanics is not well understood. We performed experimental studies and developed mathematical models to identify how reductions in turgor pressure affect anther transverse stiffness and resonant frequency. We determined that both the transverse stiffness and the resonant frequency decreased monotonically with decreasing turgor pressure. In the discussion that follows, we discuss the ecological significance of the relationship between turgor pressure and anther stiffness/ resonant frequencies and address other insights garnered from the present study.

### Importance of anther transverse stiffness and resonant frequency

Sufficiently high transverse stiffness and resonant frequencies are important for the function of poricidal anthers. Buzz pollinating bees can weigh upwards of 20 times that of the anther they are pollinating; thus, the anther deforms considerably under the mass of the bee during buzz pollination [29]. High transverse stiffness ensures that the anther itself does not experience excessive bending that may cause the pores to collapse under the weight of the bee, thus precluding the extraction of pollen. Instead, most of the deformation occurs at the filament supporting the anther, which leaves the pollen pores intact so that pollen may be ejected from the anther when buzzed.

The resonant frequency of the anther ensures that bees can remove pollen from the anther with relatively low energetic costs by leveraging vibration amplification. The mean anther resonant frequency of the freshly ablated anther was about 625 Hz in experimental studies (Fig. 7), but the weight of the bee can lower these values [29] into the approximately 100 - 400 Hz buzzing frequency range reported in buzz pollinating bees [6, 41]. After 60 minutes post-ablation, the resonant frequency decreased 20% on average. This reduction may shift the anther’s resonant frequency outside of the range that a particular bee species can buzz at, thereby increasing the energetic expense required to extract pollen or preventing pollen extraction altogether.

Although plants demonstrate incredible resilience to stressors and have mechanisms to stabilize turgor pressure (e.g., osmotic adjustment) [42], turgor pressure can be influenced by environmental factors. For example, water availability and soil moisture content directly impact a plant’s capacity to absorb water [43, 42]. Consequently, persistent drought-like conditions may reduce the anther turgor pressure, in turn reducing transverse stiffness and resonant frequencies. Other factors, such as humidity, temperature, and light intensity, can periodically affect turgor pressure throughout the day [44, 45, 46]. Each of these factors influences the evaporation phase of transpiration. As water evaporates from the anther surface, the water potential within the anther cells decrease, thereby creating a negative pressure that pulls water into the anther tissue [47]. Reductions in turgor pressure can occur if the evaporation rates exceed the rate at which the water is drawn into the tissue [43].

We hypothesize that fluctuations in turgor pressure may influence the foraging behavior of buzz-pollinating insects as they seek to exploit favorable anther mechanics. Bumblebees have been shown to adjust their buzzing frequency in response to the biophysical properties of the flowers they pollinate [48], suggesting that they can adapt to mechanical changes in poricidal anthers. However, it remains necessary to determine if or how buzz-pollinating bees adjust their buzzing characteristics or foraging behavior when environmental stressors drive changes in plant turgor pressure.

### Poroelasticity affects anther mechanics

Consistent with previous modeling efforts, we assumed that the influence of turgor pressure on the beam-like anther could be idealized as an axial tensile load [34]. The magnitude of the tensile load was proportional to the turgor pressure and anther cross sectional area. Increases in the tensile load increase anther transverse stiffness and resonant frequencies. However, this modeling approach assumes that the turgor pressure is constant and uniform throughout the anther. Bending deformations cannot cause fluid flow through the porous networks embedded within the plant tissue. In reality, anther bending generates spatiotemporal changes in turgor pressure which cause fluid transport through the porous anther tissue [49].

Our dynamic nanoindentation experiment (Fig. 8) highlights the importance of fluid flow through the poroelastic anther tissue. At low frequency excitation, the fluid has adequate time to flow between pores. At higher excitation frequencies, the timescale of bending deformations is faster than that of fluid flow. Thus, fluid does not have sufficient time to flow between pores as the anther deforms. The result is that the fluid perceptively stiffens the anther, thereby resulting in a higher effective storage modulus. Buzz pollinating insects buzz at frequencies generally ranging from 100 - 400 Hz [6], which suggests anther deformations will occur at time scales that prohibit fluid transport. Within this regime, the anther is stiffer when excited dynamically than it is when excited statically.

Therefore, there are limitations to modeling the effect of turgor pressure as a static tensile load rather than directly modeling the poroelastic behavior. First, fluid transport and its effect on anther deformations cannot be predicted. This is problematic for low-frequency excitation, where the fluid flow may considerably change the mechanical relationship between input force and output displacement of the anther. Second, the modulus must be known as a function of frequency to utilize dynamic models. Poroelastic models may account for the increase in perceived modulus via first principles, whereas this correction must be made empirically if treating the turgor pressure effectively as a static axial load. Despite its limitations, the present model is simple to implement compared to a poroelastic structural frameworks, and is well suited to predict static transverse stiffness and anther resonant frequencies. It also requires less model inputs compared to a poroelastic model, where tissue permeability and other parameters must be known. Future high-fidelity models of poricidal anthers may consider incorporating these model features.

## Author contributions

Conceptualization – MJ; Data curation – MJ, CC, MA, JM; Formal analysis – MJ, MA, JM; Funding Acquisition – MJ; Investigation – MA, JM, CC; Methodology – MJ, MA, JM, CC; Project administration – MJ; Resources – MJ, MA, CC; Software – MA, JM; Supervision – MJ; Validation – MA, JM; Visualization – MJ, MA, JM; Writing (original draft) – MJ, MA, JM; Writing (review and editing) – MJ, CC, MA, JM

## Data Availability

Processed primary data to support the findings of this study are openly available in https://zenodo.org/records/11223625.

